# Sleeve gastrectomy promotes colitis-associated colorectal cancer in a murine model via a modified gut microbiome

**DOI:** 10.1101/2022.06.04.494831

**Authors:** James N. Luo, Renuka S. Haridas, Tammy Lo, Ali Tavakkoli, James Yoo, Eric G. Sheu

## Abstract

Colorectal cancer (CRC) remains the third leading cause of cancer death in the United States with an alarming rise among young (<50-years-old) patients.^1^ Epidemiologically, obesity appears to be a risk factor for CRC.^1^ Although bariatric surgery has been shown to be associated with decreased risk for most cancers, studies to date on bariatric surgery and CRC continue to yield conflicting results.^2^ One possible explanation for this seeming irreconcilability is the inherent heterogeneity of CRC with its varied mechanisms. This is likely compounded by the differing bariatric operations currently employed. Here, we sharpen our focus and investigate how the most performed bariatric operation, sleeve gastrectomy (SG), affects colitis-associated CRC. Using a murine model, we found that SG significantly exacerbates both colitis and colitis-associated CRC. Using a germ-free (GF) microbiota transplant model, we found that the post-SG microbiota, when transplanted into GF mice, is capable of independently recapitulating the tumor-promoting phenotype of SG. Our results suggest that the postsurgical microbiome plays a key causal role in the increased risk for CRC after SG. This finding represents the first step in our understanding of this complex relationship that is at the intersection of two rising public health threats.

---

Colorectal cancer (CRC) remains the third leading cause of cancer death in the United States with an alarming rise among young (<50-years-old) patients.^1^ Epidemiologically, obesity appears to be a risk factor for CRC.^1^ Although bariatric surgery has been shown to be associated with decreased risk for most cancers, studies to date on bariatric surgery and CRC continue to yield conflicting results.^2^ One possible explanation for this seeming irreconcilability is the inherent heterogeneity of CRC with its varied mechanisms. This is likely compounded by the differing bariatric operations currently employed. Here, we sharpen our focus and investigate how the most performed bariatric operation, sleeve gastrectomy (SG), affects colitis-associated CRC.

Diet-induced-obese (DIO) mice were randomized to SG or sham surgery using techniques previously reported by our group.^3^ Following surgical recovery, mice were challenged with azoxymethane (AOM) and dextran sulfate sodium (DSS) to induce colon tumor formation (Figure 1a). Colitis severity was assessed by daily measurement of weight, diarrhea, and the presence of hematochezia during each cycle of DSS. Screening colonoscopies were performed on all mice prior to sacrifice. At sacrifice, the number of tumors in each mouse was manually counted. Stool and ceca were collected from each mouse using sterile techniques and stored for later use.

**Figure 1.**
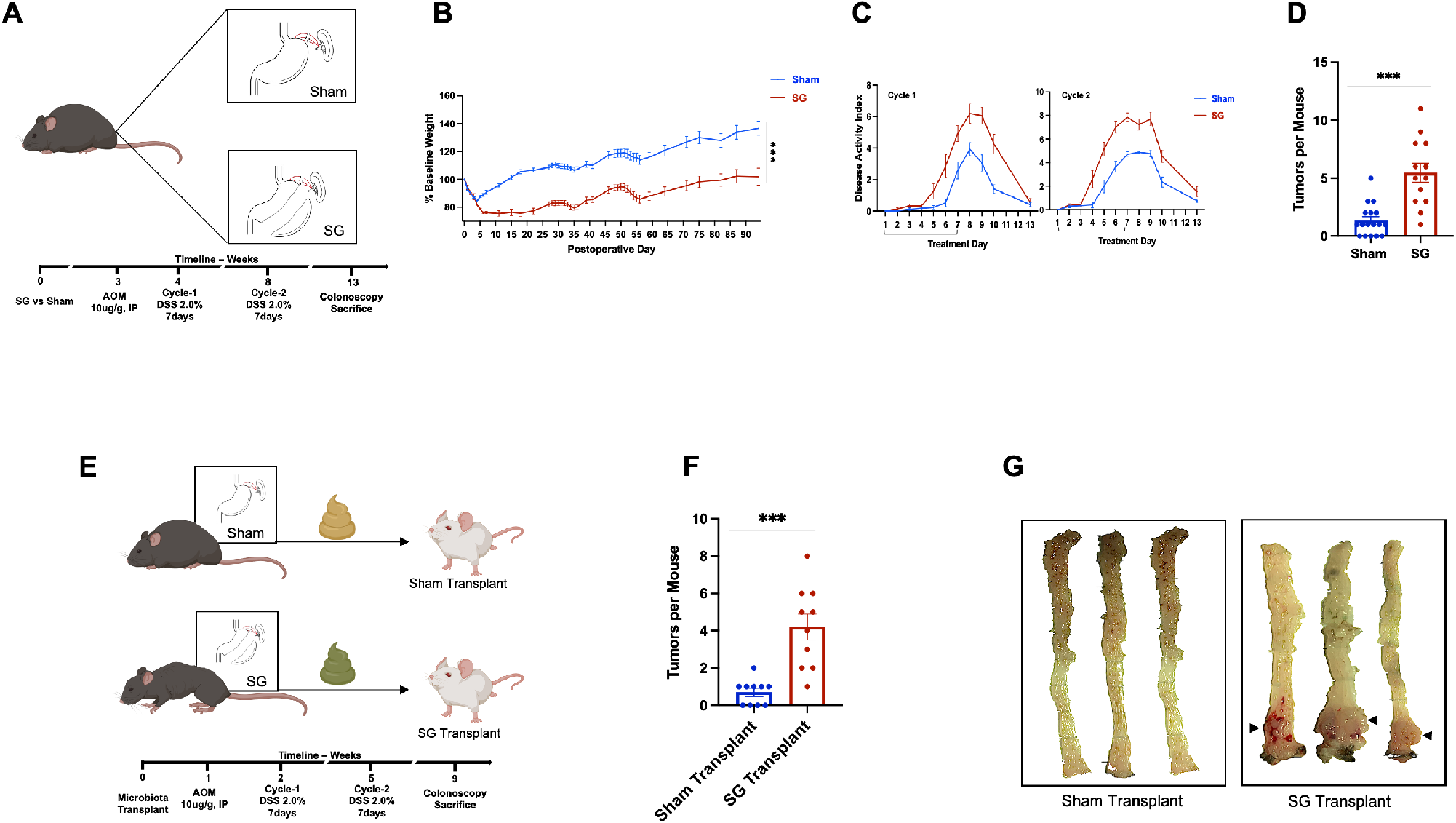
SG increases colitis-associated colonic tumors and its effect is recapitulated via microbiota transplant. (A) Experimental timeline of surgical SG and colitis-associated cancer model. Age- and weight-matched diet-induced-obese (DIO) mice were randomized to SG or sham surgery. AOM and DSS treatment began 2 weeks after surgical recovery. Animals were sacrificed 13 weeks following initial surgery. SG n=13, sham n=17. (B) SG mice experienced significantly less weight gain while fed a high-fat-diet (HFD) compared to sham mice. Data points represent mean ±SEM, ***p<0.001, two-way ANOVA. (C) SG mice exhibited significantly higher disease activity index (DAI) during both cycles, accounted by more severe diarrhea, weight loss, and hematochezia. Horizontal brackets depict days of DSS administration. Data points represent mean ±SEM, p<0.001 for both cycles, two-way ANOVA. (D) SG resulted in >4-fold increase in the average numbers of tumors per mouse. Columns represent mean ±SEM, ***p<0.001, Student’s T-test. (E) Cecal microbiota transplant (CMT) was performed by transplanting cecal microbiota from the surgical cohort into age- and weight-matched germ-free mice via gavage. Following engraftment, recipient mice were challenged with the same AOM/DSS regimen as their surgical donors; n=10 per group (F) Ex-GF mice conventionalized with SG donors (SG-R) developed nearly 5-fold increase in tumors per mouse than those with sham donors (sham-R). Columns represent mean ±SEM, ***p<0.001, Student’s T-test. (G) Representative gross images of the colon from CMT-recipient mice. Arrow heads indicate location of the greatest tumor density (distal colon).

As expected, SG mice exhibited significantly less weight gain on high-fat-diet (HFD) compared to sham mice (p<0.001, Figure 1b)^6^. During each cycle of DSS, SG mice showed significantly worsened colitis as evidenced by increased diarrhea and hematochezia, in addition to weight loss (p<0.001 for each cycle, Figure 1c). Moreover, SG mice experienced >4-fold increase in the average number of tumors per mouse (1.4 vs. 5.5, p<0.001, Figure 1 d). Analysis of colonic tissue cytokine expression showed SG mice with significantly higher levels of several pro-inflammatory cytokines, including TNF-a, IL-1ß, IL-6, Il-23 (Supplementary Figure 1a).

Among the potential biologic links between bariatric surgery and CRC, the gut microbiome has emerged as a leading candidate. Numerous previous studies have described the enduring and conserved shifts in the microbiome following bariatric surgery.^3^ Similarly, an accumulating body of evidence has linked the microbiome with CRC pathogenesis.^4^ We thus elected to further investigate this potential mechanistic axis. We began with 16S rRNA sequencing of stool collected at sacrifice. We found that among the top microbiome changes in the SG cohort were blooms in the relative abundance of *Akkermansia* and *Bacteroides* (Supplementary Figure 1b). *Akkermansia* enrichment is a consistently reported finding following bariatric surgery, however, recent reports have linked it to worsening colitis.^7^ These observations prompted us to hypothesize that the post-SG microbiome could be contributing to tumor development.

To interrogate whether the microbiome played a causal role in the SG tumor-promoting phenotype, we transplanted the cecal microbiota from our surgical cohort into germ-free mice of the same genetic background (Figure 1 e). These recipient mice were similarly fed a HFD and challenged with the same AOM/DSS regimen as their surgical donors. Remarkably, recipients of the SG microbiota (SG Transplant) experienced a nearly 5-fold increase in the average number of tumors per mouse compared to recipients of the sham microbiota (Sham Transplant) (0.7 vs 4.2, p<0.001, Figure 1f). Grossly, tumor burden is concentrated in the distal rectum (Figure 1g). The observed tumor phenotypes in the recipient mice virtually recapitulated those observed in their respective donors, suggesting that the microbiome plays a causal role in SG-mediated tumorigenesis in colitis-associated CRC.

To our knowledge, this is the first study directly linking bariatric surgery and the post-bariatric microbiome with CRC development in a preclinical model. In this work, we purposely narrowed our scope by studying the effects of the most performed bariatric operation on a specific CRC subtype. This focused approach allowed us to identify key phenotypic and mechanistic effects of SG in cancer development. Several groups have recently reported a potential association between bariatric surgery and worsening inflammatory bowel disease.^5, 6^ However, none has yet to definitively show this link extending to cancer.

The precise mechanism(s) relating the post-SG microbiome with colitis-associated CRC remains elusive. One of the hallmark microbial changes following bariatric surgery has been enrichment of *Akkermansia,* which has long been considered a metabolically favorable shift.^7^ However, several recent works have suggested that increased abundance of *Akkermansia* results in worsening colitis, possibly due to its mucolytic activity on colonic epithelial barrier function.^8, 9^ It is conceivable that this diminished barrier protection resulting from *Akkermansia-* derived mucolytic activity exposes the colonic epithelium to greater luminal carcinogen, although this currently remains speculative. Another potential explanation for our observation is SG’s impact on bile acid signaling. Prior works have shown increased expression of the G-protein-coupled bile acid receptor, TGR5, in colitis-associated cancer tissues. ^10^ Our group has previously shown increased TGR5 agonism following SG via altered bile acid signaling,^3^ thus suggesting another possible link between bariatric surgery and CRC.

Our work is limited by the stochastic nature of our model. Both SG and AOM/DSS can exert profound effects on the gut microbiome. When combining these effects into a single model, it is difficult to precisely isolate the microbial element responsible for our phenotype. Nonetheless, we believe that given the lack of robust studies in this area, our work represents an important first-step in elucidating this complex biology. This work also reveals several logical next steps for future studies. Does Roux-en-Y gastric bypass (RYGB) exert a similar effect as SG? How do other types of CRC behave in response to bariatric surgery? These questions will further advance our understanding of this complex biology. Lastly, the findings herein are preclinical and validation in humans will be needed before they can be successfully translated into clinical practice.

In summary, using a murine model, we showed SG promotes colitis-associated CRC, and that microbiota transplant is sufficient to independently recreate this phenotype. Our results suggest that the postsurgical microbiome plays a key causal role in the increased risk for CRC after SG. This finding represents the first step in our understanding of this complex relationship that is at the intersection of two rising public health threats.

## Supplementary Methods

### Mice

All animals were age-, weight-, and sex-matched for all experiments. For surgery, 11 week-old, diet-induced-obese (DIO), male, C57BL/6J mice were purchased from Jackson Laboratory (Bar Harbor, ME). Animals were housed under standard conditions of climate-controlled environment with 12-hour light-and-dark cycle. Mice were maintained on high-fat-diet (HFD, 60% Kcal fat; RD12492; Research Diets Inc., NJ). On arrival, mice were acclimated for at least 1-week prior to any experiments. For cecal microbiota transplant (CMT), male germ-free (GF) C57BL/6 mice were bred internally by the Massachusetts Host-Microbiome Center (https://researchcores.partners.org/cctm/about) and maintained there in gnotobiotic isolators. All experiments were approved by the Institutional Animal Care and Use Committee of Brigham and Women’s Hospital.

### Surgical Model

Sleeve gastrectomy (SG) surgical model has been previously described by our group.^1^ Briefly, a 1.5cm midline laparotomy incision was made under general anesthesia via isoflurane. The short gastric arteries were ligated to fully mobilized the stomach. A greater curve based gastric sleeve was constructed by removing 80% of the glandular and 100% of the non-glandular stomach via stapler. Sham operation involved the identical midline incision and ligation of short gastric arteries. Postoperatively, mice were recovered on Recovery Gel Diet (Clear H2O, Westbrook, ME) for 6 days before resuming HFD, and maintained on it for the duration of the experiment. SG cohort n=13; sham cohort n=17.

### Colitis-associated Cancer Model

Following surgery, mice were allowed 2-weeks to recover from procedural trauma. Colitis-associated cancer was induced using protocol described by Snider and colleagues with the following modifications.^2^ Azoxymethane (AOM, Milipore Sigma cat-A5486) was reconstituted using sterile deionized water and mice received a 1-time intraperitoneal injection (10μg/g) 2-weeks followings surgery. 1-week later, mice underwent 2 cycles (7-days per cycle) of dextran sulfate sodium (DSS, MP Biomedicals cat-160110) at 2% concentration. Mice were given a 2-week recovery period between DSS cycles. Weight, stool consistency, and hematochezia presence were tracked daily. Disease activity index (DAI) was calculated using the system described by Park and colleagues.^3^ 13-weeks after surgery, screening colonoscopy was performed on all mice prior to sacrifice. At sacrifice, the entire colon was removed, irrigated, and opened. Number of tumors were counted. For analytic consistency, only macroscopically visible tumors were counted. ~30% of the colonic tissue was then excised and fixed in formalin for histology, and the remainder frozen in liquid nitrogen and stored in −80°C for later use. Stool was collected and immediately frozen in liquid nitrogen and then stored in −80°C for later use. Cecum was resected, immediately frozen in liquid nitrogen, and then stored in −80°C for microbiota transplant.

### Cecal Microbiota Transplant (CMT)

Frozen ceca harvested at sacrificed was thawed in an anaerobic environment. Ceca were suspended in PBS with 0.05% cysteine (C-6852, Sigma-Aldrich). Pooled (n=4) SG or sham ceca were homogenized. The resultant slurry was centrifuged at 700g for 1 minute. Supernatant was collected and used for inoculation. All steps performed were under strict anaerobic condition. Weight-matched 8-week-old male GFC57Bl/6 mice were inoculated with 200μL of the inoculum via gavage. For the entire duration of the experiment, mice were maintained in gnotobiotic isolators at the Massachusetts Host-Microbiome Center under 12-hour light and dark cycle with a constant temperature (21°C ± 1°) and humidity (55–65%). Starting 1-week posttransplant, recipient mice underwent the same AOM/DSS regimen as described above. This timeline allowed for 2-weeks following transplant before introducing DSS, thus ensuring stable engraftment. At the conclusion of the experiment, mice were sacrificed in the same manner as described above.

### mRNA Cytokine Expression

mRNA expression was measured using techniques previously described by our group.^4^ Briefly, RNA was isolated from distal (~1cm from the anal verge) colonic mucosa using RNeasy mini kits (Qiagen, Germany). Superscript III First-Stand Synthesis System (Life Technologies, CA) was used for cDNA construction. SYNR green reporter was used for RT-PCR. TATA-box-binding protein (Tbp) was used for as reference gene.^5^ Results were analyzed using the comparative delta-delta Ct method.

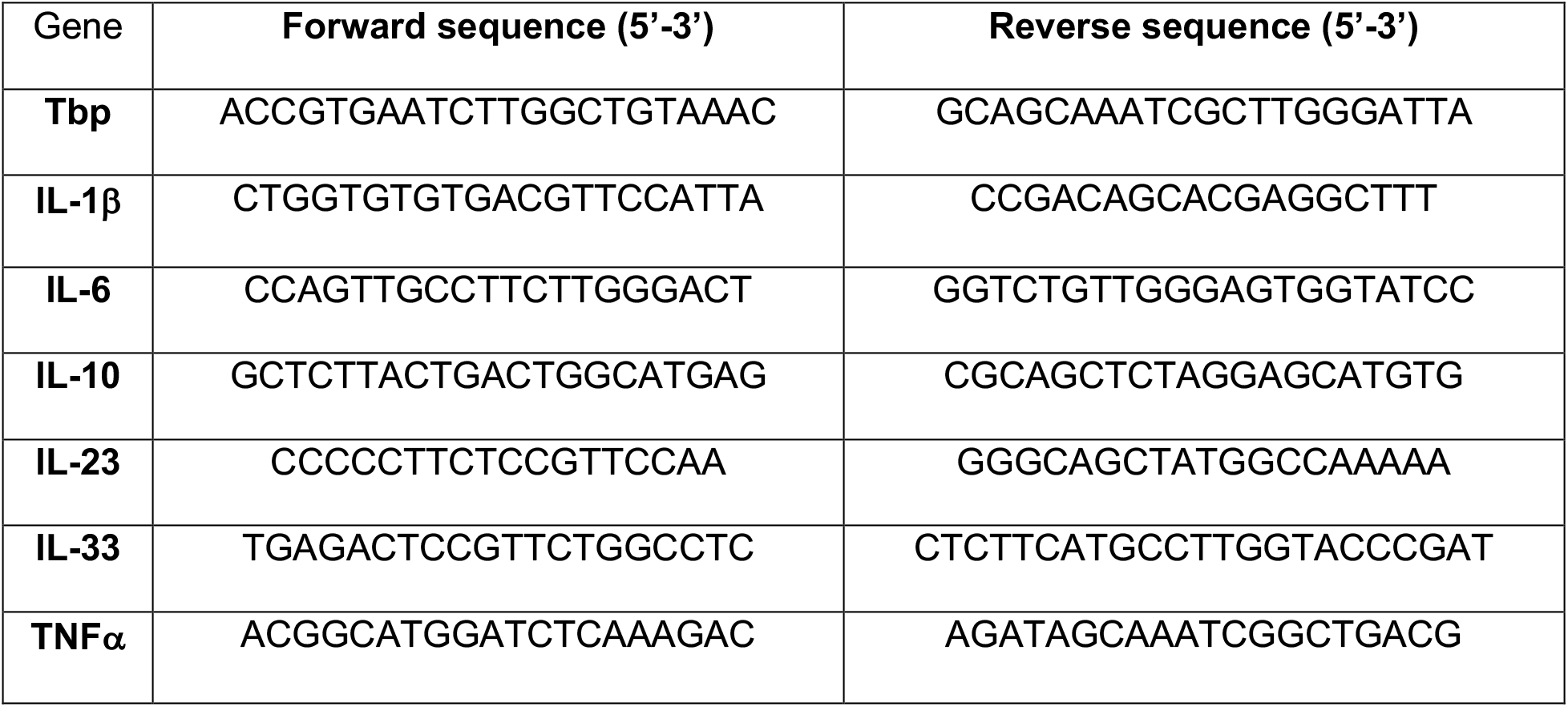

### 16S rRNA Sequencing

Stool samples were submitted to the Massachusetts Host-Microbiome Center for DNA extraction and amplification. Fecal microbiota DNA was extracted using QIAamp Fast DNA Stool Mini Kit (Qiagen, Germany). Variable region 4 (v4) of the 16S rRNA gene was amplified on the Illumina MiSeq platform. All microbiome analyses were done by an independent third party, Harvard Chan Microbiome Analysis (HCMA) Core (https://hcmph.sph.harvard.edu/hcmac/). Detailed analytic workflow can be obtained via HCMA website. Briefly, FASTQ sequences were resolved using the paired-end DADA2 pipeline. Reads were filtered for quality control using a MAXEE score of 1. Filtered reads were used to generate the OTUs. UPARSE OTU analysis pipeline was used for OTU clustering and taxonomy prediction with percent identity 0.97 and minimum cluster size of 2. The SILVA database was used for taxonomy prediction. All resultant plots were generated using R version 3.6.2.

### Statistical Analysis

Data were plotted and analyzed using GraphPad Prism 7 (GraphPad Software, San Diego, CA). Statistical significance was determined using two-way ANOVA, Welch’s t-test, and Student’s t-test wherever appropriate.

## Acknowledgement

Elements of the figures created with BioRender.com.

---

**Supplementary Figure 1.**
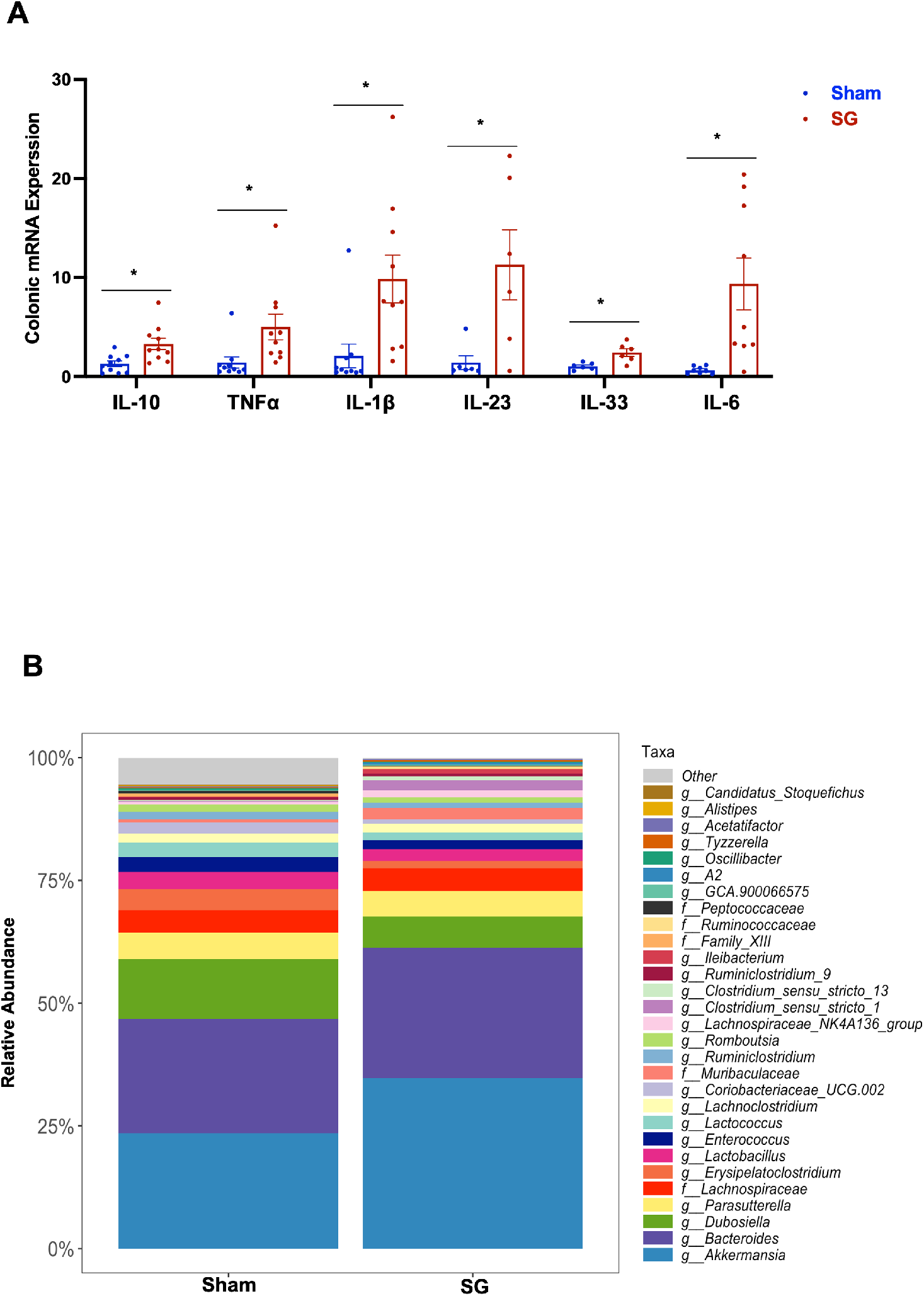
(A) Cytokine gene expression of key inflammatory cytokines in the distal colonic mucosa of the surgical cohort; n=6-10 per group, *p<0.05, Student’s T-test. (B) SG was associated with changes in the relative abundance of several key microbial species, including a marked increase in *Akkermansia* and *Bacteroides*.

